# In situ evidence that mast cells release mutant NLRP1 in keratoacanthomas from multiple self-healing palmoplantar carcinoma

**DOI:** 10.1101/2025.09.12.675637

**Authors:** Alexandra Dobre, Tudor E. Fertig, Andrei M. Niculae, Adelina M. Cohn, Antoanela Curici, Razvan T. Andrei, Daciana S. Marta, Victor E. Peteu, Roua G. Popescu, George C. Marinescu, Gabriela Turcu, Ana M. Forsea, Daniela A. Ion, Mihaela Gherghiceanu, Roxana I. Nedelcu

**Affiliations:** Carol Davila University of Medicine and Pharmacy, Bucharest, Romania; Elias University Emergency Hospital, Dermatology, Bucharest, Romania; Victor Babeş National Institute of Pathology, Bucharest, Romania; Pathology Department, “Dr. Carol Davila” Central Military Emergency University Hospital, Bucharest, Romania; Laboratoire National De Santé, Dudelange, Luxembourg; Synevo Romania, Bucharest, Romania; Colentina Clinical Hospital, Bucharest, Romania; Faculty of Medicine, Titu Maiorescu University, Bucharest, Romania; Asociaţia Independent Research, Bucharest, Romania; Blue Screen SRL, Bucharest, Romania; Derma 360 Clinic, Bucharest, Romania; Health and Medical University Potsdam, Germany

## Abstract

NLRP1 is an inflammasome sensor protein expressed in barrier tissues of humans. Its activation in response to microbes or cellular stress triggers a cascade of molecular events, leading up to IL1β-driven inflammation and pyroptosis. Rare germline mutations of NLRP1 cause its persistent activation, resulting in autoinflammatory syndromes. Multiple self-healing palmoplantar carcinoma (MSPC) is one such syndrome, characterized by the appearance of recurrent keratoacanthomas (KAs) on the palms and soles. Here, we aimed to compare the subcellular localization of mutant NLRP1 in MSPC-associated lesions, to wild-type NLRP1 in non-MSPC KAs and in skin from healthy donors. Using mass spectrometry, immunohistochemistry and immunoelectron tomography, we found that NLRP1 localized to mast cell (MC) granules in all samples, a novel finding which implicates MCs in NLRP1-associated responses in human skin. Moreover, we found that MCs expressing the A66V pathogenic variant of NLRP1 overpopulated MSPC-KAs, infiltrated the epidermis and degranulated, a behavior not seen in other lesions from this study. The released granules had the highest NLRP1 protein content and also contained NLRP3 and IL1β, indicating coexistence of inflammasome pathways within MCs. Taken together, our findings establish cutaneous MCs as a NLRP1 reservoir in health and disease, opening a new area of research in NLRP1-related syndromes.

## 1. INTRODUCTION

NLRP1 (NLR family pyrin domain containing 1) is a cytosolic sensor protein with a major role in innate immunity. In response to pathogens or other stress signals, NLRP1 can participate in the formation of an intracellular protein complex termed the “inflammasome” [1]. This assembly involves the recruitment of the adapter protein ASC (apoptosis-associated speck-like protein containing a CARD), which facilitates the activation of caspase-1. Once activated, caspase-1 mediates the maturation and secretion of pro-inflammatory cytokines IL-1β and IL-18 and the induction of pyroptosis [2].

In humans, NLRP1 pathogenic variants can lead to a variety of clinical syndromes depending on the affected protein domain, mainly involving the skin, cornea and lungs [3-6]. Of these, multiple self-healing palmoplantar carcinoma (MSPC) is characterized by recurrent keratoacanthomas (KAs) that develop on palmoplantar skin and sporadically on other epithelia lacking hair follicles [7]. Responsible are either one of three germline heterozygous mutations (A54T, A66V, M77V) affecting the pyrin domain (PYD) of NLRP1, causing spontaneous self-oligomerization of the protein, aberrant inflammasome activation and subsequent proliferation of select keratinocytes [7].

Keratinocytes are known as the main NLRP1 reservoir in human skin [7-10], however granulocytes, lymphocytes, macrophages and dendritic cells also express NLRP1[11]. RNA-seq data from the Human Protein Atlas indicates basophils as having the highest potential for NLRP1 protein expression among immune cells [12,13].

Mast cells (MCs) are tissue-resident immune cells of hematopoietic origin, which differentiate from a basophil-MC common progenitor. Although not yet described to contain NLRP1, MCs contain NLRP3 within secretory granules and can produce IL1β following NLRP3 inflammasome activation [14,15]. Mutations of NLRP3 cause it to become constitutively activated in MCs, initiating overproduction of IL1β. This contributes to a spectrum of clinical entities named cryopyrin-associated periodic syndromes (CAPS), in which the skin, joints and central nervous system can be affected [14,16].

We report for the first time that skin MCs harbor NLRP1 within granules. We further show massive and specific degranulation of MCs in different KAs of a MSPC patient with NLRP1^A66V^ and suggest that release of granules containing this hyperactive NLRP1 variant contributes to the pathogenesis of KAs in this syndrome.

## 2. RESULTS

### 2.1. Different skin lesions from a MSPC patient have divergent patterns of inflammasome activation

A 49-year-old male with a history of multiple central hyperkeratotic lesions on the palms and soles (**Figure 1a**) in the context of MSPC with NLRP1^A66V^, presented with symmetrically distributed and well-defined erythematous-squamous plaques, some with erosions and crusting. These initially appeared on the dorsal feet and later spread to the extremities, trunk and head (**Figure 1b**). An infection was ruled out because of unresponsiveness to antibiotics/antifungals and negative PAS staining. Histopathology suggested psoriasis based on the presence of psoriasiform hyperplasia, parakeratosis containing some neutrophiles, thinning of the suprapapillary plates and focal hypogranulosis (**Figure 1c and inset**). However, an overlap with allergic contact dermatitis was suspected due to (a) atypical development and clinical presentation of plaques, (b) positive patch-tests to lanolin alcohol and nickel(II) sulfate hexahydrate, (c) good response of lesions to oral corticotherapy and lanolin-free moisturizers, and (d) lack of recurrence after stopping corticotherapy.

**Figure 1.**
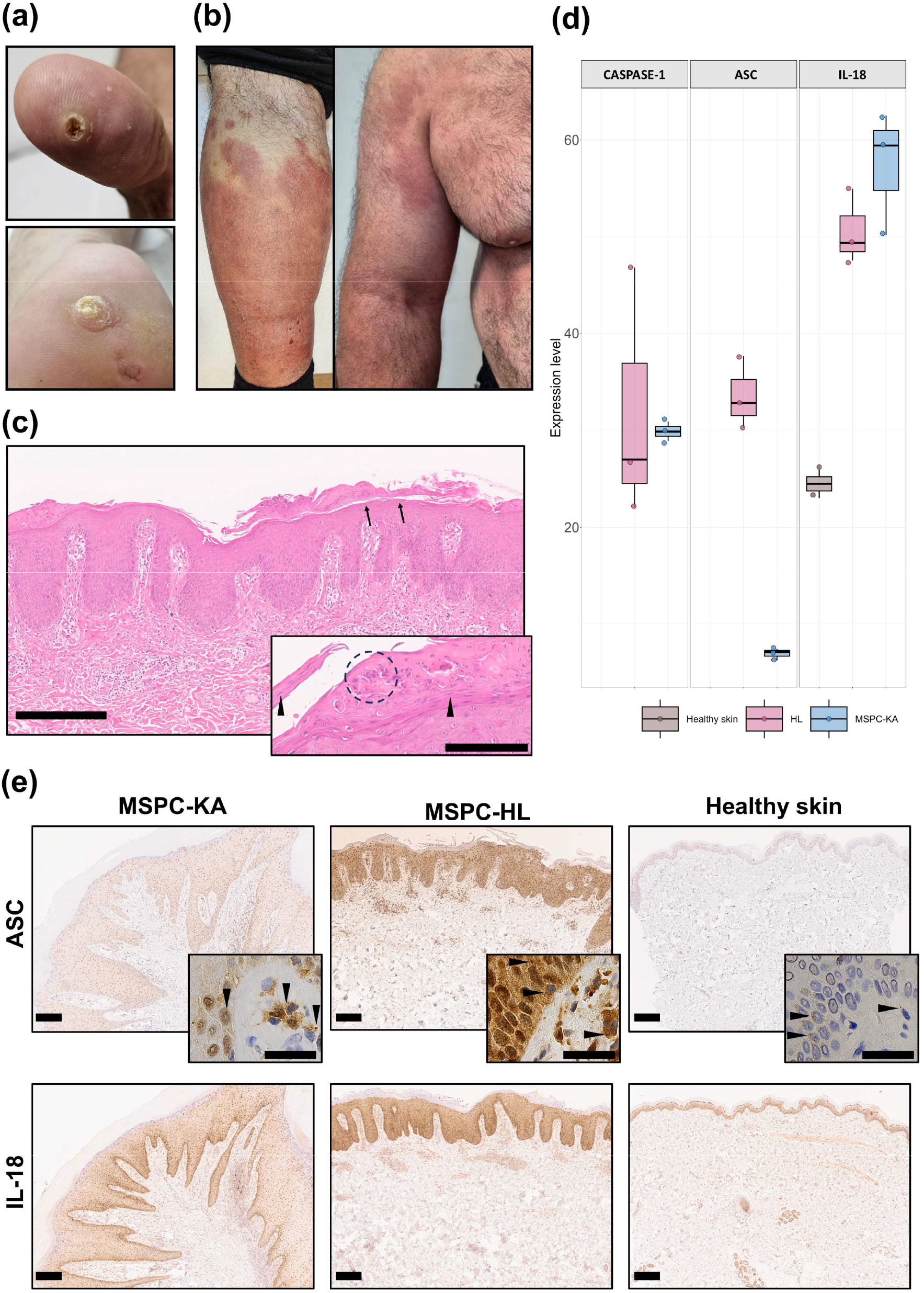
Inflammasome activation is different between two types of skin lesions from a patient with multiple self-healing palmoplantar carcinoma (MSPC). (**a**) The MSPC patient presented keratoacanthomas (KAs) at different stages of evolution on the palmar and plantar surfaces; (**b**) The MSPC patient suddenly developed a widespread eruption of well-defined erythemato-squamous plaques; (**c**) Hematoxylin and eosin staining (HE, 10x and 20x inset) of an abdominal lesion from the MSPC patient, showed psoriasiform acanthosis with elongation of the rete ridges, thinned suprapapillary plates, hypervascular dermal papillae, neutrophil clusters (circle) in the parakeratotic stratum corneum (arrows heads) and hypogranulosis (arrows); Scale bars are 300 and 100 µm for inset; (**d**) Mass spectrometry (MS) showing higher protein expression levels for caspase-1, apoptosis-associated speck-like protein containing a CARD (ASC) and interleukin-18 (IL-18) the abdominal hyperkeratotic inflammatory lesion (HL) and MSPC-KAs, compared to healthy skin. ASC levels were highest in MSPC-HL, but were below detection levels in healthy skin; (**e**) Spatial distribution of inflammasome-associated proteins ASC (top row and insets) and IL-18 (bottom row) in MSPC lesions and healthy skin. The epidermis of MSPC-HL was strongly positive for ASC and IL-18, which appeared uniformly distributed throughout the cell layers. In MSPC-KAs, both proteins localized to cell clusters in the dermis, but epidermal staining was weaker when compared to MSPC-HL. Granular material suggestive of ASC speck formation is indicated by arrow heads in insets. Scale bars are 200 µm for all images in (e).

MSPC patients characteristically develop palmoplantar KAs and it remains unclear why KAs fail to form on other anatomical regions. The sudden appearance of these extensive atypical hyperkeratotic inflammatory lesions (HL) in our MSPC patient, presented a unique opportunity to compare their lesional microenvironment to that of different plantar KAs from the same individual (biopsied before, concomitantly and after MSPC-HL diagnosis), in an attempt to isolate the etiology of MSPC-KAs.

We first asked whether there are differences in inflammasome activation between the two lesion types and we employed MS and IHC to investigate protein expression. As expected, both MSPC-KAs and MSPC-HL showed higher abundance of key inflammasome-associated proteins, compared to healthy skin (**Figure 1d, e**). MS analyses revealed that whereas caspase-1 and IL-18 had similar levels of expression in the two MSPC lesions, ASC was significantly overexpressed in MSPC-HL (**Figure 1d**). This may indicate higher priming of inflammasome pathways and/or accumulation and persistence of stable ASC-based supramolecular complexes in MSPC-HL tissue.

These complexes (known as “specks”) are a hallmark of inflammasome activation and are readily detectable by light microscopy as discrete puncta in the nucleus and cytosol [17; 18]. Consistent with MS data, the strongest immunoreactivity for ASC was in the MSPC-HL epidermis. Here, and to a lesser extent in the subjacent dermis, cells exhibited a dispersed punctate pattern indicative of different stages of speck assembly (**Figure 1e, top row and inset**). In MSPC-KA lesions, ASC also had a strong, granular signal associated with dermal cell clusters, contrasting with an unexpectedly weak staining of keratinocytes in the overlying epidermis (**Figure 1e, top row and inset**). The differences in staining intensity for IL-18 were less striking, but still favored the epidermis of MSPC-HL, where it appeared uniformly distributed across cell layers. In MSPC-KAs, IL-18 also concentrated to cell clusters in the dermis and in the basal layer of the epidermis (**Figure 1e, bottom row**). These results demonstrated the existence of different hotspots for inflammasome activation in the two MSPC lesions, likely involving site-specific skin cell populations and pathogenic mechanisms.

### 2.2 MCs accumulate in MSPC-KAs

Given that both MSPC-KAs and MSPC-HL showed increased inflammasome activity but divergent ASC staining patterns, we hypothesized that a cell-specific distribution of NLRP1^A66V^ may at least in part explain the formation of characteristic KAs on palmoplantar skin, but not on skin from other anatomical regions.

In IHC, NLRP1 distribution and staining intensity mimicked our spatial data on ASC. Keratinocytes from MSPC-KAs were positive for NLRP1, with a nucleo-cytoplasmic staining pattern and the intensity gradually decreasing towards the upper epidermal layers (**Figure 2a**). Contrary to previous reports [7], the NLRP1 signal was more intense in the dermis than the epidermis, with specific cell populations presenting a granular distribution within the cytoplasm (**Figure 2a, top row inset**). Inversely, NLRP1^A66V^ expression was highest in the epidermis of the MSPC-HL sample, but comparatively reduced in the dermis, mainly clustering around blood vessels (**Figure 2a**). Healthy skin showed the least NLRP1 staining intensity in both epidermis and dermis (**Figure 2a**), although the granular distribution persisted in cells surrounding blood vessels. Positivity of immune cell granules for NLRP1 was confirmed on a human lymph node specimen (**Supplementary Figure S1a**), whereas the isotype control did not bind (**Figure 2a**).

**Figure 2.**
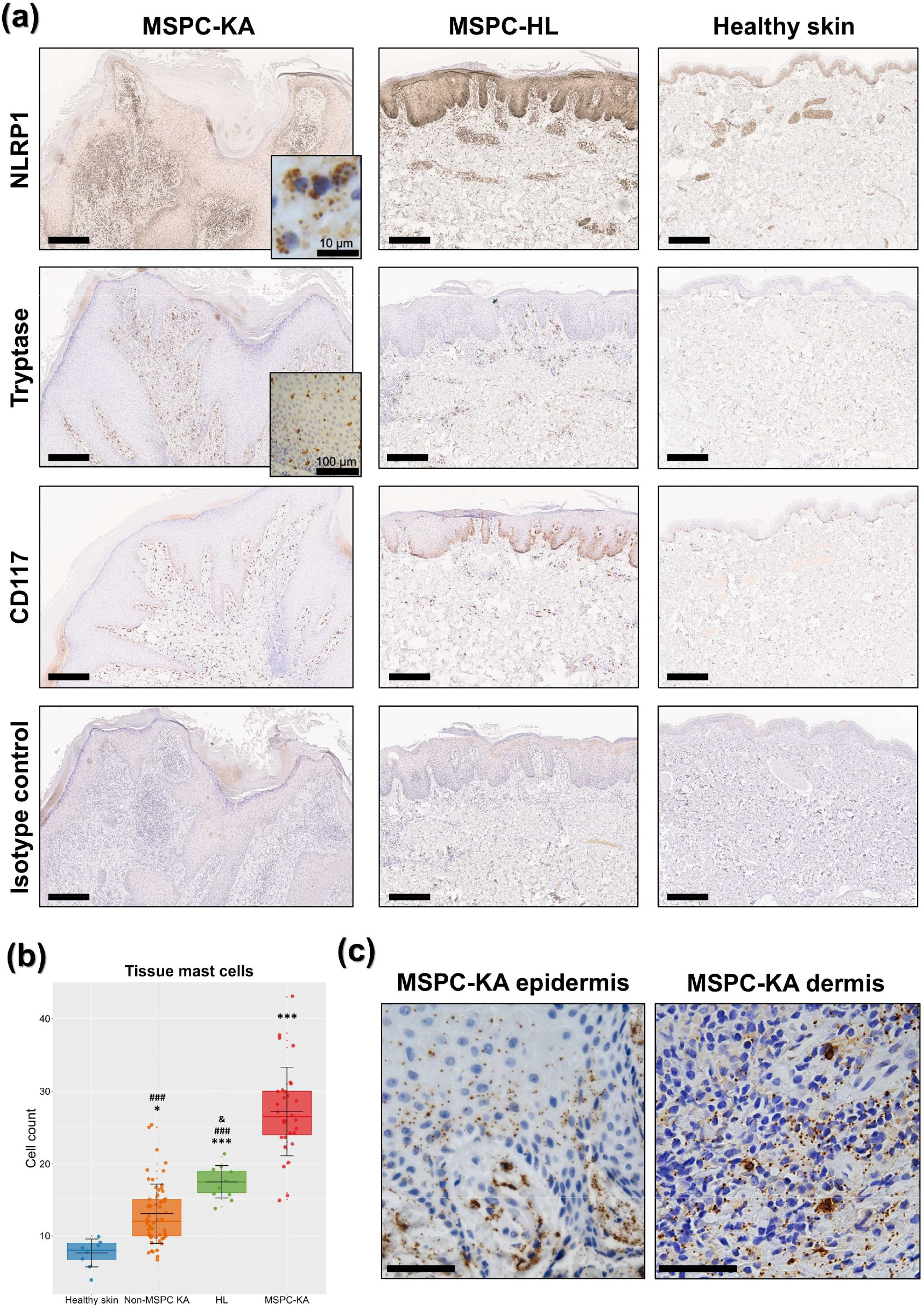
Immunohistochemistry of keratoacanthomas (KAs) in multiple self-healing palmoplantar carcinoma (MSPC). (**a**) In MSPC-KAs, NLRP1^A66V^ was more highly expressed in the dermis and was concentrated in cell granules (top row inset, scale bar = 10 µm). In an atypical hyperkeratotic inflammatory lesion (HL), NLRP1^A66V^ was overrepresented in the epidermis. CD117 and tryptase staining demonstrated more mast cells (MCs) in MSPC-KAs, including epidermal infiltration (second row inset, scale bar = 100 µm). The isotype control did not bind. Scale bars = 300 µm. (**b**) Quantification of MCs in the analysed samples, presented as means ± standard deviation (SD). Statistical significance is shown against healthy skin—*, MSPC-KA group—#, and the non-MSPC KA group—&, where */& p <.05; ***/### p <.001. (**c**) MCs in MSPC-KAs were degranulated, with granular material dispersed in epidermis and dermis. Scale bars = 50 µm.

To investigate whether some of the numerous NLRP1-positive cells in the MSPC-KA dermis were MCs, we stained using tryptase and CD117/c-kit (**Figure 2a**). Indeed, KA samples from the MSPC patient had significant MC accumulation (27.2/mm2), above MSPC-HL (17.5/mm2) or healthy skin (7.6/mm2) (**Figure 2b**). In MSPC-KAs but not MSPC-HL, MCs frequently infiltrated epidermal layers (**Figure 2a, second row inset**).

Under higher magnification we further observed that MCs in MSPC-KAs, but not MSPC-HL, were degranulated. Tryptase-positive granular material was widely distributed within connective tissue and taken up in the epidermis, up to the spinous layer (**Figure 2c**). To verify if these are common features of all KAs, we used tryptase staining on six KAs from patients not diagnosed with MSPC (**Supplementary Figure S1b**). As expected [19], these presented MCs in higher numbers (13/mm2) than healthy skin, however below counts in MSPC-KAs and even MSPC-HL (**Figure 2b**). Importantly, we could not identify MC epidermal infiltration, nor MC degranulation, indicating their distinct contribution in MSPC-KA pathogenesis.

The MSPC patient was negative for D816V KIT in whole blood and had normal serum tryptase (<11.4 µg/L), limiting the possibility of MC neoplastic transformation.

Taken together, these results suggest that plantar KAs from a MSPC patient with the NLRP1^A66V^ pathogenic variant have a specific cellular landscape, dominated by high numbers of degranulating MCs, which can infiltrate the epidermis.

### 2.3 NLRP1 localizes to MC granules

For more precise subcellular localization of NLRP1 we employed immunoelectron microscopy (IEM). Morphologically, most MCs from MSPC-KA were degranulated, ranging from having empty granules and granules fused with the plasma membrane, to complete fragmentation of cells, with release of granular complexes in the extracellular space (**Figure 3a**). By contrast, MCs from MSPC-HL and healthy skin had typical appearance (intact granule architecture, presence of plasma membrane folds) and localized in proximity of blood vessels (**Figure 3b, c**).

**Figure 3.**
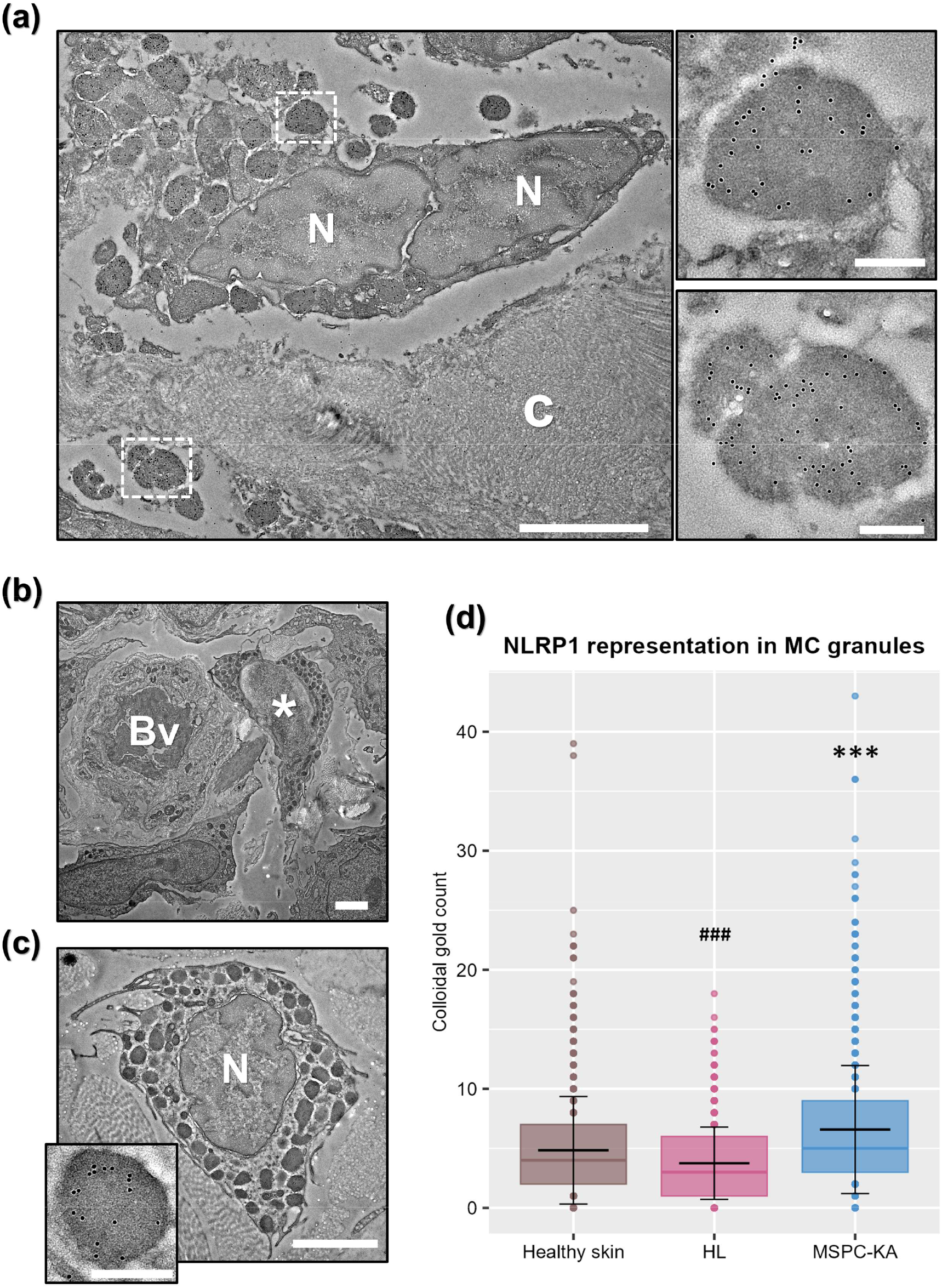
NLRP1 is ubiquitous in skin mast cell (MC) granules. (**a**) MCs in keratoacanthomas (KAs) from a multiple self-healing palmoplantar carcinoma (MSPC) patient were degranulated. Scale bar = 2 µm. c - collagen, N - nucleus. MC granules in MSPC-KA presented NLRP1^A66V^ accumulation, as exemplified by higher magnifications of boxed areas. Gold labeling is seen as black dots. Scale bars = 250 nm. (**b**) MCs from an atypical hyperkeratotic inflammatory lesion (HL) were morphologically normal (*) and clustered around blood vessels (Bv). Scale bar = 2 µm. (**c**) Normal MC in healthy skin presenting NLRP1 in granules (inset, scale bar = 250 nm). Scale bar = 2 µm. (**d**) Quantification of colloidal gold demonstrated MCs in MSPC-KA had more NLRP1 (means ± standard deviation). Statistical significance is shown against healthy skin—*, and MSPC-KA—#, where ***/### p <.001.

Using two different, commercially available anti-NLRP1 antibodies, we demonstrated that NLRP1 accumulates in granules of MCs from both lesional and healthy skin, regardless of granule morphology and degranulation status of the cell, indicating NLRP1 is a ubiquitous component of skin MCs (**Figure 3a, c and inset**). By contrast, an isotype control did not bind (**Supplementary Figure S2a**). Next, we asked whether there are differences in MC NLRP1 content between samples, so we manually counted and averaged colloidal gold per granule, under identical immunolabeling conditions. In this context, this method of quantification is more precise than western blotting of tissue lysates, as it measures protein content in the subcellular compartment of interest, unbiased by overall representation of MCs in tissue. We found that NLRP1^A66V^ from MSPC-KA was overrepresented compared to NLRP1^A66V^ in MSPC-HL or wild-type NLRP1 in healthy skin (**Figure 3d**), further suggesting MCs may serve as a reservoir of hyper-reactive NLRP1^A66V^ during MSPC-KA pathogenesis.

In accordance with IHC, NLRP1 was localized in keratinocytes in all samples, however it unexpectedly clustered around intermediate filaments and desmosomes (**Supplementary Figure S2b**).

### 2.4 NLRP1 and NLRP3 can colocalize in MC granules

MCs are known to contain all components of the NLRP3 inflammasome and to secrete IL1β in a NLRP3-dependent manner [14,15]. Given our finding that NLRP1^A66V^ was present in MC granules from the MSPC patient, we were curious whether it spatially overlapped with NLRP3 and if it could therefore share elements of the inflammasome machinery during activation. First, we labeled MSPC-KA against pro-IL1β/IL1β (hereafter IL1β) and found it well represented in both intracellular MC granules and in MC-derived extracellular granular material (**Figure 4a and inset**). This demonstrated that MCs in MSPC-KA secrete this pro-inflammatory cytokine in the extracellular space during degranulation.

**Figure 4.**
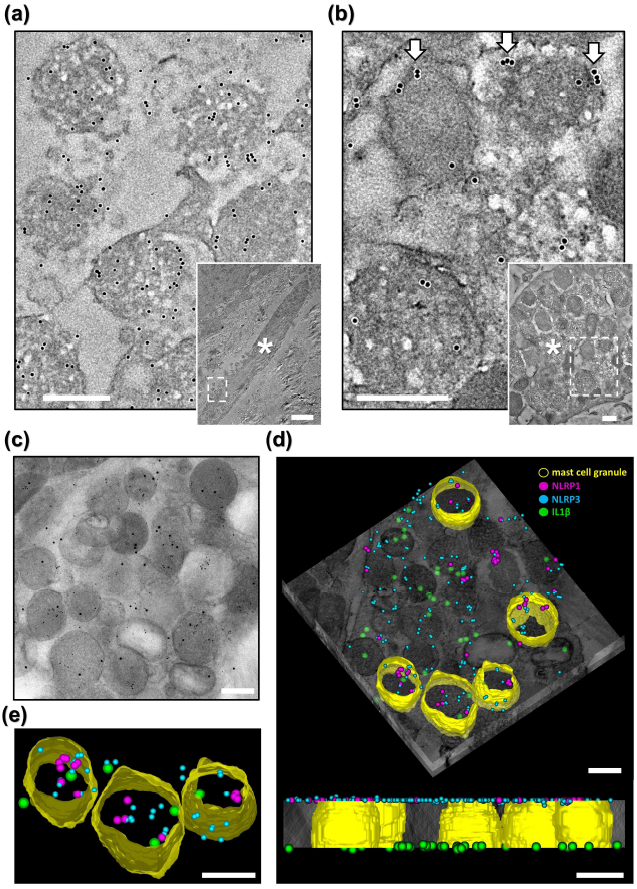
NLRP1^A66V^, NLRP3 and IL1β colocalize to mast cell (MC) granules in a MSPC-keratoacanthoma. (**a**) High magnification of box in inset, confirming pro-IL1β/IL1β in granular material released by a MC (*). (**b**) High magnification of box in inset, showing MCs (*) contain NLRP3, which frequently localized to the periphery of granules (arrows). (**c**) Projection from tomographic tilt series, showing colocalization of gold labels (NLRP1^A66V^, NLRP3, IL1β) in MC granules. (**d, top**) Reconstructed tomogram with superimposed contours of five MC granules (yellow), presenting colocalization of NLRP1^A66V^ (magenta), NLRP3 (turquoise) and IL1β (green). (**d, bottom**) Side-view showing distribution of labels to each side of the imaged resin section. (**e**) Detail of segmented MC granules. Scale bar for inset (a) = 2.5 µm, all others = 300 nm.

Next, we confirmed that some MC granules also contained NLRP3 (**Figure 4b and inset**) and released it during degranulation. NLRP3 was frequently localized at the periphery of cytoplasmic granules and at the plasma membrane in degranulating MCs (**Figure 4b**), in agreement with a recently published study showing a role for NLRP3 in granule exocytosis [20]. Electron tomography on double-labeled sections revealed that NLRP1 and NLRP3 can colocalize to some, but not all MC granules (**Supplementary Figure S2c, d, Supplementary Video S1**), raising the possibility of compartmentalization of inflammasome pathways within MCs. Finally, triple-labeling revealed that both sensor proteins can colocalize with IL1β, in both intra- and extracellular MC granules (**Figure 4c-e, Supplementary Video S2**), confirming that multiple inflammasome sensor proteins can simultaneously inhabit MC granules and may act in tandem during activation of MCs in MSPC-KAs.

## 3. DISCUSSION

Here, we demonstrated that skin MCs can act as a reservoir for NLRP1, a critical but lesser known inflammasome sensor protein. Although novel, this finding is not completely surprising, as MCs express pattern recognition receptors and harbor all components of the NLRP3 inflammasome pathway [15], some of which are also required for NLRP1 inflammasome assembly.

We next showed that MCs are directly involved in the pathogenesis of KAs in a MSPC patient with NLRP1^A66V^. In three different plantar lesions, separated from initial diagnosis by 1, 8 and 10 years, MCs infiltrated the epidermis and NLRP1^A66V^ was released in the dermis and epidermis through widespread MC degranulation. These features were conspicuously absent in a HL specimen from the abdominal skin of the same MSPC patient or in KAs from 6 non-MSPC patients.

The activation of MCs in MSPC-KAs may be a response to NLRP1-inflammasome activation in local keratinocytes, which would then release pro-inflammatory factors. However, this is difficult to reconcile with two key findings: (1) that the highest intensity of ASC and NLRP1 staining were in the MSPC-HL epidermis, where keratinocytes also express the hyperactive NLRP1^A66V^, yet do not form KAs, and (2) that compared to healthy skin, MSPC-HL presented high numbers of MCs prone to activation by keratinocyte inflammatory cues, but these MCs had normal morphology.

Alternatively, the particular MC behaviour in the plantar skin of this MSPC patient may be the result of intrinsic NLRP1-inflammasome activation in response to yet unknown triggers. Activation of select MC clusters and release of granular material would induce keratinocyte inflammatory responses, increasing MC recruitment within the lesion.

In support of this, we showed that MC granular material spread throughout the basal and spinous layers of MSPC-KAs. These granules had higher NLRP1 content than equivalents in MSPC-HL and healthy skin and also harbored NLRP3 and IL1β. The latter can be converted to its active form within intact granules via the NLRP1 and NLRP3 inflammasome pathways, but also in the extracellular environment by endogenous proteases [20,21], thereby driving keratinocyte proliferation [21]. Additionally, tryptase released during MC degranulation can cleave protease activated receptor 2 (PAR-2) on keratinocytes [22], stimulating non-specific particle uptake through phagocytosis [23]. In MSPC-KAs, this would allow enrichment of keratinocytes with components of the NLRP1/NLRP3 inflammasome pathways and other granule components known to stimulate keratinocyte proliferation (e.g. histamine, serotonin and keratinocyte growth factor) [24,25]. The pro-inflammatory activity of extracellular sensor proteins has been previously demonstrated for ASC and mutant NLRP3 oligomers. After being internalized by macrophages through phagocytosis, these complexes can induce ASC aggregation, caspase-1 activation and IL-1β release [26]. The characteristic palmoplantar epidermal hyperplasia in MSPC-KAs may therefore be conditioned by aberrant, NLRP1-dependent activity of tissue-resident MCs.

This study has two important limitations. First, in absence of functional studies, we cannot definitively ascertain whether MC degranulation is a prerequisite for MSPC-KA formation, or if it is a local response mechanism which merely modulates lesion development. As murine NLRP1 lacks the PYD domain [7], an in vitro experiment would be required, involving co-culture of human keratinocytes with MCs expressing mutant NLRP1. Second, because we could only include a single MSPC patient, carrying the NLRP1^A66V^ pathogenic variant, we are unable to extrapolate our findings to other MSPC variants. This is further hindered by the rarity of these diseases, with only 5 MSPC families known worldwide [27].

In summary, we showed for the first time that NLRP1 is a component of skin MC granules and can colocalize with NLRP3 and IL1β, indicating that MCs have more avenues for inflammasome activation than previously known. We also showed that when MCs from plantar skin express NLRP1^A66V^, they proliferate, infiltrate the epidermis and degranulate, contributing to the pathogenesis of KAs in this variant of MSPC. This opens a new area of research in NLRP1-related syndromes and, if confirmed in other patients, would translate clinically to accessible therapies targeting MC activity.

## 4. MATERIALS AND METHODS

### 4.1. Patients and samples

All subjects gave written informed consent prior to biopsy. The MSPC patient included here was diagnosed and genetically characterized in a previous study [7]. Four samples were collected from this patient: three different KAs from plantar regions (in 2016, 2023 and 2025) and a sample from an atypical hyperkeratotic inflammatory lesion (HL) on the abdomen (in 2023). Biopsies were collected from the elbow region of two clinically healthy volunteers. The study included 6 archived formalin-fixed paraffin-embedded (FFPE) KA specimens from different patients with no clinical history of MSPC. Human lymph node and tonsil FFPE specimens were included as NLRP1 and ASC positive controls. This study was performed in conformity with the Declaration of Helsinki and received approval from the ethics committees at Victor Babeş National Institute of Pathology and Elias University Emergency Hospital, Bucharest, Romania.

### 4.2. Light microscopy and immunohistochemistry

Hematoxylin-eosin was done by routine protocol. Immunohistochemistry (IHC) with the primary antibody of interest (**Supplementary Table S1**) was done using an Autostainer Link 48, following manufacturer specifications (Dako, Agilent Technology, US). An isotype control and omission of primary antibody were used to verify non-specific binding. Digital slides were generated using an automated slide scanner (Aperio AT2, Leica Biosystems, US). Images were processed by Contrast Limited Adaptive Histogram Equalization (CLAHE) in Fiji [28], using the same settings for each type of staining.

### 4.3. Mass spectrometry

Biopsies were collected in cold PBS, then 50 mg from each sample was disrupted in a Mini-Beadbeater (Biospec, US). Samples were then centrifuged for 20 minutes at 18,000 g and the protein concentration from the supernatant was quantified by Bradford assay. A total of 50 μg of protein containing supernatant was diluted with urea buffer to the final volume of 40 μL, followed by the addition of 25 μL of 100 mM DTT and incubation at room temperature for 60 minutes. The sample was then alkylated with 26.25 μL of 300 mM iodoacetamide and kept in the dark at room temperature for an additional 90 minutes followed by a quenching reaction with 50 μL of DTT 100mM at room temperature for 60 minutes. The reaction mix was finally diluted to 500 μL with 50 mM ammonium bicarbonate, followed by addition of trypsin (1 μg /μL; Trypsin Gold, V528A, Promega, US) at a 1:50 enzyme-to-substrate ratio, with incubation at 37°C for 16 hours. The reaction was stopped by adding 3 μL of formic acid. Peptides were cleaned up using solid phase extraction (MonoSpin C18, GL Sciences, Japan), dried at speed-vac and rehydrated to 2 μg/μL using 5% ACN + 0.1% HCOOH. Peptide concentration was measured using NanoDrop 1000 (Thermo Fisher Scientific, US). From each sample, 5 μg peptides were loaded into nanoACQUITY UPLC system (Waters, US) and separated using an Eksigent 5C18-CL-120 (300 μM ID, 150 mm length) column coupled with a TripleTOF 5600+ mass spectrometer (both AB Sciex, USA). Peptide samples (5 μg) were injected using Solvent A (0.1% formic acid) and separated on a 5-90% gradient Solvent B (0.1% formic acid in acetonitrile) over 90 minutes, at a flow rate of 5 μL/min, with the column maintained at 55°C. Each sample underwent three replicate runs. Mass spectrometric (MS) analysis was carried out using electrospray ionization in positive mode, with a spray voltage of 5500 V and a source temperature of 200°C. Data acquisition was performed on a TripleTOF 5600+ system in DIA SWATH-MS mode, with 64 variable isolation windows. The MS1 survey scans covered an m/z range of 400–1250, while MS2 spectra were collected in high-sensitivity mode from 100 to 2000 m/z. The accumulation period was set to 0.049 s, and ions were scanned within 55 ms windows in high-sensitivity mode, giving a total cycle time of 3.5 s. Peptide and protein identification were performed using PeakView 2.2, Skyline 22.2.0.255 and DIA-NN 1.9.

### 4.4. Immunoelectron microscopy

Biopsies were fixed in a solution comprising 4% paraformaldehyde, 0.5% glutaraldehyde, and 1.4% sucrose (pH 7.4), then embedded in London Resin White (LR White, Agar Scientific, UK). Ultra-thin sections were mounted on formvar and carbon-coated nickel grids, then incubated with the primary antibody of interest (**Supplementary Table S1**) and a gold-conjugated secondary antibody at 1:50 dilution (Aurion, The Netherlands). Negative controls had primary antibodies omitted and an isotype control was used to exclude nonspecific binding to MC Fc receptors. A blocking solution for goat antibodies was used to reduce nonspecific binding (Aurion, The Netherlands). Imaging was done using a 4×4k Ceta camera, on a Talos 200C transmission electron microscope (Thermo Fisher Scientific, US).

### 4.5. Immunoelectron tomography

LR White sections (300 nm) were adhered to nickel grids precoated with 0.01% poly-L-lysine and without formvar or carbon. For double-labeling, grids were incubated on each side, consecutively, with two primary antibodies raised in different species (mouse and rabbit, **Supplementary Table S1**). For triple-labeling, grids were incubated on one side with a mixture of two antibodies raised in different species (mouse and rabbit, **Supplementary Table S1**), then either a rabbit or mouse primary antibody was used on the other side. Secondary antibodies had different colloidal gold diameters for each target (6, 10 and 15 nm, Aurion, The Netherlands). Absence of cross-contamination between section faces was verified by immunolabeling control grids on one side, while the other side was incubated with PBS. Acquisition of single axis tilt series and tomographical reconstruction and segmentation in IMOD-eTomo [29], were done as described elsewhere [30].

### 4.6. Data analyses

Quantification of MCs was performed on tissue sections stained for tryptase. For each case, tryptase-positive cells were manually counted in ten consecutive high-power fields (HPF, 40× objective), starting from a hotspot. Each experimental group included approximately 27 observations. To quantify the NLRP1 content of MC granules, cells were first identified on randomly selected ultrathin sections from the MSPC and healthy skin specimens, immunolabeled under identical conditions. Entire MCs were imaged, then colloidal gold was counted on all granules of each cell. Each experimental group included approximately 1,058 observations.

Statistical analyses were performed using one-way ANOVA, followed by Tukey’s post hoc test for pairwise comparisons between groups. Statistical significance is presented only for comparisons with p-values <.05. The post hoc statistical power for the applied tests was 1. Graphical representations were generated using R 4.4.1.

## SUPPLEMENTARY

**Video S1. NLRP1 and NLRP3 colocalize to MC granules in MSPC-KA**. In situ tomogram and segmentation showing colocalization of NLRP1 (magenta) and NLRP3 (green) to MC granules in MSPC-KA (related to Figure S3c,d).

**Video S2. NLRP1, NLRP3 and IL1β colocalize to MC granules in MSPC-KA**. In situ tomogram and segmentation showing colocalization of NLRP1 (magenta), NLRP3 (turquoise) and IL1β (green) to MC granules in MSPC-KA (related to Figure 4c-e). for (a-b).

---

**Supplementary Figure S1.**
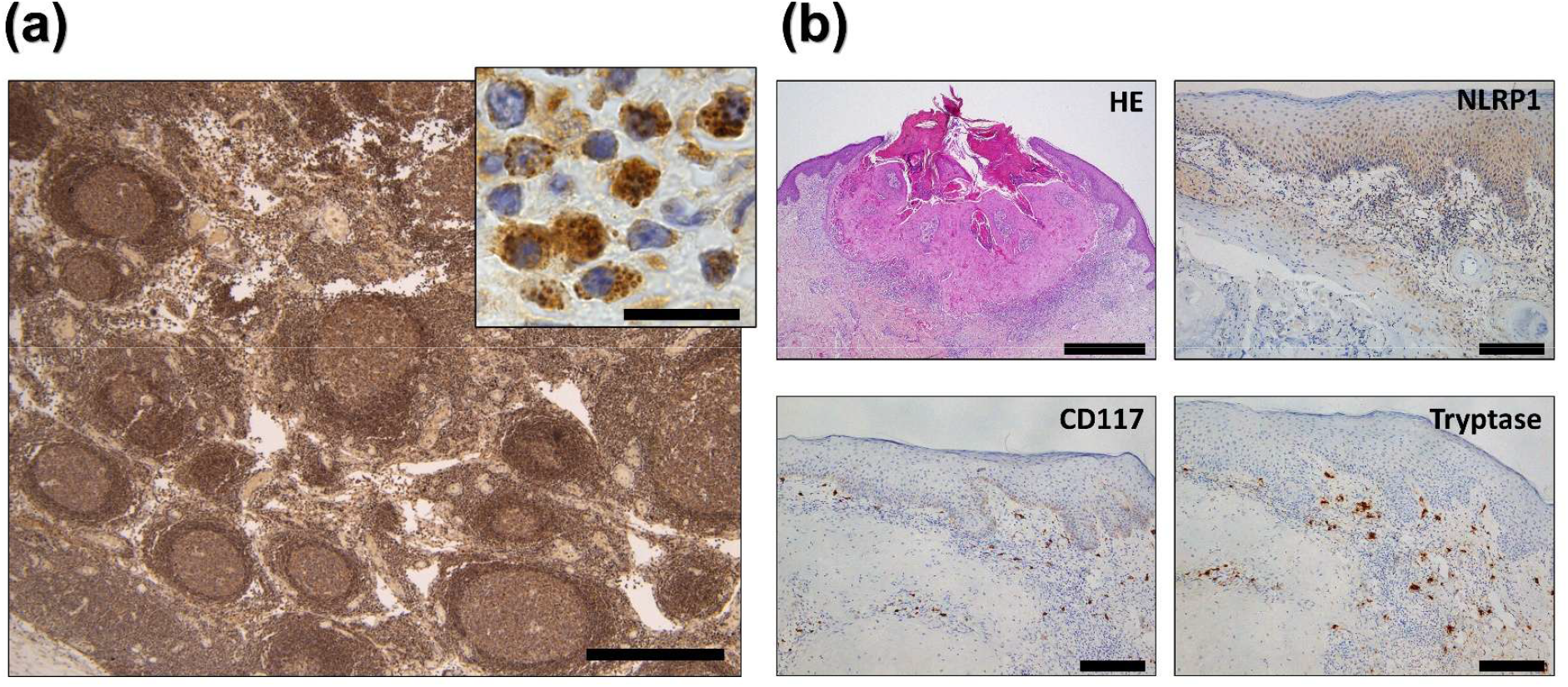
Additional immunohistochemistry controls for NLRP1 localization and for mast cell involvement in KA lesions. **(a)** Human lymph node stained against NLRP1, showing highest proportion of positive cells in the mantle zone of lymphoid follicles. Scale bar = 500 µm. Higher magnification **(inset**, scale bar = 20 µm) demonstrates localization of NLRP1 to granules of immune cells in lymph node. **(b)** Representative regions from non-MSPC KA, under different stains, showing absence of mast cell degranulation and epidermal infiltration. Scale bars are: 500 and 20 µm for (a) and inset, respectively; 1 mm for HE staining in (b) and 200 µm for all others panels in (b).

**Supplementary Figure S2.**
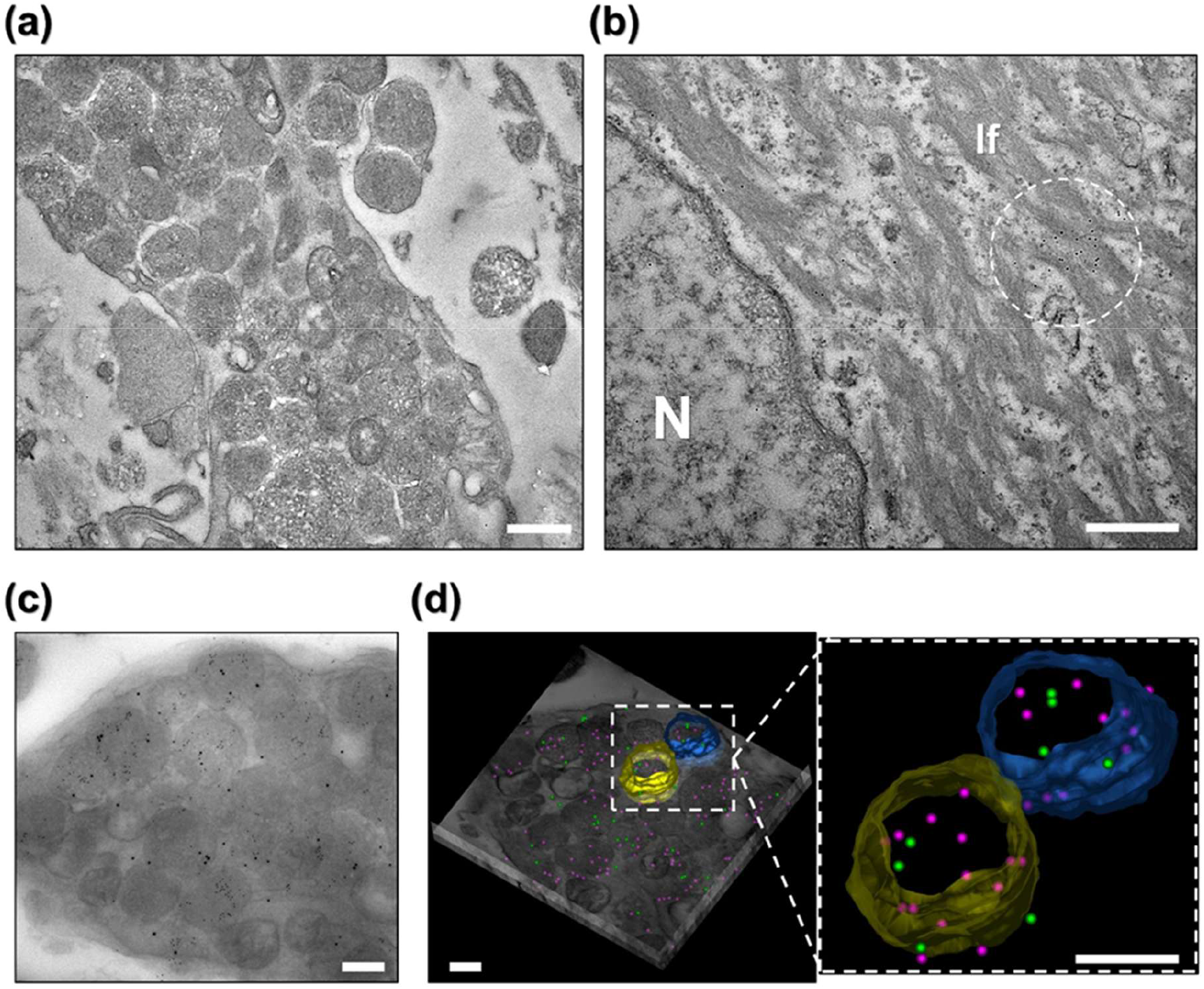
Immunoelectron microscopy negative control and immunoelectron tomography of NLRP1 and NLRP3 colocalization in mast cell granules. **(a)** A rabbit isotype control did not significantly bind to mast cell (MC) granules. **(b)** NLRP1 localized to intermediate filaments (If) of keratinocytes. Representative accumulation of colloidal gold can be seen as black dots (circled area). *N - nucleus*. **(c)** Projection from tomographic tilt series, showing colocalization of 10 and 15 nm gold particles (NLRP1^A66V^ and NLRP3, respectively) in granules from a MC in MSPC-KA. **(d)** Reconstructed tomogram with superimposed drawn contours of two MC granules (yellow and blue), presenting colocalization of NLRP1^A66V^ (magenta) and NLRP3 (green). Detail of segmented MC granules demonstrating colocalization of NLRP1^A66V^ (magenta) and NLRP3 (green) are presented in the boxed area. Scale bars = 500 nm.

## REFERENCES

1. Martinon F, Burns K, Tschopp J. The Inflammasome: A molecular platform triggering activation of inflammatory caspases and processing of proIL-β. Mol Cell [Internet]. 2002 [cited 2025 Jun 21];10:417–26. Available from: https://pubmed.ncbi.nlm.nih.gov/12191486/

2. Bauernfried S, Hornung V. Human NLRP1: From the shadows to center stage. Journal of Experimental Medicine [Internet]. 2021 [cited 2025 Jun 21];219. Available from: 10.1084/jem.20211405

3. Barry K, Murphy C, Mansell A. NLRP1-A CINDERELLA STORY: a perspective of recent advances in NLRP1 and the questions they raise. Commun Biol [Internet]. 2023 [cited 2025 Jun 21];6:1–7. Available from: https://www.nature.com/articles/s42003-023-05684-3

4. Mitchell PS, Sandstrom A, Vance RE. The NLRP1 inflammasome: new mechanistic insights and unresolved mysteries. Curr Opin Immunol [Internet]. 2019 [cited 2025 Jun 21];60:37–45. Available from: https://pubmed.ncbi.nlm.nih.gov/31121538/

5. Burian M, Schmidt MF, Yazdi AS. The NLRP1 inflammasome in skin diseases. Front Immunol. 2023;14:1111611.

6. Calabrese L, Fiocco Z, Mellett M, Aoki R, Rubegni P, et al. Role of the NLRP1 inflammasome in skin cancer and inflammatory skin diseases. British Journal of Dermatology [Internet]. 2024 [cited 2025 Jun 21];190:305–15. Available from: https://pubmed.ncbi.nlm.nih.gov/37889986/

7. Zhong FL, Mamaï, M, Sborgi L, Saad A, Hiller S, et al. Germline NLRP1 Mutations Cause Skin Inflammatory and Cancer Susceptibility Syndromes via Inflammasome Activation. Cell [Internet]. 2016 [cited 2024 Apr 18];167:187–202. Available from: 10.1016/j.cell.2016.09.001

8. Sand J, Haertel E, Biedermann T, Contassot E, Reichmann E, et al. Expression of inflammasome proteins and inflammasome activation occurs in human, but not in murine keratinocytes. Cell Death Dis [Internet]. 2018 [cited 2025 Jun 21];9:1–14. Available from: https://www.nature.com/articles/s41419-017-0009-4

9. Feldmeyer L, Keller M, Niklaus G, Hohl D, Werner S, Beer HD. The Inflammasome Mediates UVB-Induced Activation and Secretion of Interleukin-1β by Keratinocytes. Current Biology [Internet]. 2007 [cited 2025 Jun 21];17:1140–5. Available from: https://www.sciencedirect.com/science/article/pii/S0960982207015084

10. Marie J, Kovacs D, Pain C, Jouary T, Cota C, et al. Inflammasome activation and vitiligo/nonsegmental vitiligo progression. Br J Dermatol. 2014 Apr;170(4):816-23. doi: 10.1111/bjd.12691. PMID: 24734946.

11. Kummer JA, Broekhuizen R, Everett H, Agostini L, Kuijk L, et al. Inflammasome components NALP 1 and 3 show distinct but separate expression profiles in human tissues suggesting a site-specific role in the inflammatory response. Journal of Histochemistry and Cytochemistry [Internet]. 2007 [cited 2025 Jun 21];55:443–52. Available from: https://pubmed.ncbi.nlm.nih.gov/17164409/

12. Karlsson M, Zhang C, Méar L, Zhong W, Digre A, et al. A single–cell type transcriptomics map of human tissues. Sci Adv [Internet]. 2021 [cited 2025 Jun 21];7. Available from: https://pubmed.ncbi.nlm.nih.gov/34321199/

13. The Human Protein Atlas [Internet]. [cited 2025 Jun 21]. Available from: https://www.proteinatlas.org

14. Nakamura Y, Kambe N, Saito M, Nishikomori R, Kim YG, et al. Mast cells mediate neutrophil recruitment and vascular leakage through the NLRP3 inflammasome in histamine-independent urticaria. Journal of Experimental Medicine [Internet]. 2009 [cited 2025 Jul 16];206:1037–46. Available from: https://pubmed.ncbi.nlm.nih.gov/19364881/

15. Bonnekoh H, Scheffel J, Kambe N, Krause K. The role of mast cells in autoinflammation. Immunol Rev [Internet]. 2018 [cited 2025 Apr 22];282:265–75. Available from: https://pubmed.ncbi.nlm.nih.gov/29431217/

16. Nakamura Y, Franchi L, Kambe N, Meng G, Strober W, Núñez G. Critical Role for Mast Cells in Interleukin-1β-Driven Skin Inflammation Associated with an Activating Mutation in the Nlrp3 Protein. Immunity [Internet]. 2012 [cited 2025 Jun 21];37:85–95. Available from: https://pubmed.ncbi.nlm.nih.gov/22819042/

17. Smatlik N, Drexler SK, Burian M, Röcken M, Yazdi AS. ASC Speck Formation after Inflammasome Activation in Primary Human Keratinocytes. Oxid Med Cell Longev. 2021 Nov 5;2021:7914829. doi: 10.1155/2021/7914829. PMID: 34777694; PMCID: PMC8589508.

18. Bryan NB, Dorfleutner A, Rojanasakul Y, Stehlik C. Activation of inflammasomes requires intracellular redistribution of the apoptotic speck-like protein containing a caspase recruitment domain. J Immunol. 2009 Mar 1;182(5):3173–82. doi: 10.4049/jimmunol.0802367. PMID: 19234215; PMCID: PMC2652671.

19. Aguiar ALM, Martins CJ, Meuser-Batista M, Carvalho V, Barreto EO, et al. A case of keratoacanthoma centrifugum marginatum with a curious mast cell accumulation at tumour sites [28]. Journal of the European Academy of Dermatology and Venereology [Internet]. 2007 [cited 2025 Apr 26];21:429–31. Available from: https://pubmed.ncbi.nlm.nih.gov/17309493/

20. Mencarelli, A., Bist, P., Choi, H.W. et al. Anaphylactic degranulation by mast cells requires the mobilization of inflammasome components. Nat Immunol 25, 693–702 (2024). 10.1038/s41590-024-01788-y

21. Macleod T, Berekmeri A, Bridgewood C, Stacey M, McGonagle D, Wittmann M. The Immunological Impact of IL-1 Family Cytokines on the Epidermal Barrier. Front Immunol [Internet]. 2021 [cited 2025 Jul 16];12:808012. Available from: https://www.frontiersin.org

22. Santulli RJ, Derian CK, Darrow AL, Tomko KA, Eckardt AJ, et al. Evidence for the presence of a protease-activated receptor distinct from the thrombin receptor in human keratinocytes. Proc Natl Acad Sci U S A [Internet]. 1995 [cited 2025 Jun 21];92:9151–5. Available from: /doi/pdf/10.1073/pnas.92.20.9151?download=true

23. Benito-Martínez S, Salavessa L, Raposo G, Marks MS, Delevoye C. Melanin Transfer and Fate within Keratinocytes in Human Skin Pigmentation. Integr Comp Biol [Internet]. 2021 [cited 2025 Jun 21];61:1546–55. Available from: https://pubmed.ncbi.nlm.nih.gov/34021340/

24. Maurer M, Opitz M, Henz BM, Paus R. The mast cell products histamine and serotonin stimulate and TNF-α inhibits the proliferation of murine epidermal keratinocytes in situ. J Dermatol Sci [Internet]. 1997 [cited 2025 Jun 21];16:79–84. Available from: https://pubmed.ncbi.nlm.nih.gov/9438912/

25. Cho KA, Kim HJ, Kim YH, Park M, Woo SY. Dexamethasone Promotes Keratinocyte Proliferation by Triggering Keratinocyte Growth Factor in Mast Cells. Int Arch Allergy Immunol [Internet]. 2019 [cited 2025 Jun 21];179:53–61. Available from: https://pubmed.ncbi.nlm.nih.gov/30909282/

26. Baroja-Mazo, A., Martín-Sánchez, F., Gomez, A. et al. The NLRP3 inflammasome is released as a particulate danger signal that amplifies the inflammatory response. Nat Immunol 15, 738–748 (2014). 10.1038/ni.2919

27. Dobre A, Nedelcu RI, Turcu G, Brinzea A, Struna I, et al. Multiple Keratoacanthomas Associated with Genetic Syndromes: Narrative Review and Proposal of a Diagnostic Algorithm. Am J Clin Dermatol [Internet]. 2024 [cited 2025 Apr 19];26:45. Available from: https://pmc.ncbi.nlm.nih.gov/articles/PMC11742465/

28. Schindelin J, Arganda-Carreras I, Frise E, Kaynig V, Longair M, Pietzsch T, et al. Fiji: An open-source platform for biological-image analysis. Nat Methods [Internet]. 2012 [cited 2025 Jul 16];9:676– 82. Available from: https://www.nature.com/articles/nmeth.2019

29. Kremer JR, Mastronarde DN, McIntosh JR. Computer visualization of three-dimensional image data using IMOD. Journal of Structural Biology. 1996 Jan-Feb;116(1):71-76. DOI: 10.1006/jsbi.1996.0013. PMID: 8742726.

30. Ceafalan LC, Fertig TE, Gheorghe TC, Hinescu ME, Popescu BO, et al. Age-related ultrastructural changes of the basement membrane in the mouse blood-brain barrier. J Cell Mol Med [Internet]. 2019 [cited 2025 Jul 16];23:819–27. Available from: https://pubmed.ncbi.nlm.nih.gov/30450815/

